# Small Extracellular Vesicles Secreted by Region-specific Astrocytes Ameliorate the Mitochondrial Function in a Cellular Model of Parkinson’s Disease

**DOI:** 10.1101/2021.04.23.441135

**Authors:** Loredana Leggio, Francesca L’Episcopo, Andrea Magrì, María José Ulloa-Navas, Greta Paternò, Silvia Vivarelli, Carlos A. P. Bastos, Cataldo Tirolo, Nunzio Testa, Salvatore Caniglia, Pierpaolo Risiglione, Fabrizio Pappalardo, Nuno Faria, Luca Peruzzotti-Jametti, Stefano Pluchino, José Manuel García-Verdugo, Angela Messina, Bianca Marchetti, Nunzio Iraci

**Affiliations:** Department of Biomedical and Biotechnological Sciences, University of Catania, Catania, 95123, Italy; Oasi Research Institute-IRCCS, Troina, 94018, Italy; Department of Biological, Geological and Environmental Sciences, University of Catania, Catania, 95125, Italy; Laboratory of Compared Neurobiology, University of Valencia-CIBERNED, Paterna, 46980, Spain; Department of Veterinary Medicine, University of Cambridge, Cambridge, CB3 0ES, United Kingdom; Department of Clinical Neurosciences, University of Cambridge, Cambridge, CB2 0QQ, United Kingdom

**Keywords:** Astrocytes, Extracellular vesicles, Exosomes, Mitochondria, High-Resolution Respirometry, Parkinson’s disease

## Abstract

Extracellular vesicles (EVs) are emerging as powerful players in cell-to-cell communication both in health and diseased brain. In Parkinson’s disease (PD) – characterized by selective dopaminergic (DAergic) neuron death in ventral midbrain (VMB) and degeneration of DAergic terminals in striatum (STR) – astrocytes (AS) exert dual harmful/protective functions. When activated by chemokine CCL3, AS promote a robust DAergic neuroprotection both in cellular and pre-clinical models of PD, with mechanisms not fully elucidated. Here we used a combination of techniques to characterize AS-EVs derived from VMB and STR, and investigated their potential to exert neuroprotection. First, we show that: (i) AS of both regions secrete small EVs of ~100 nm; (ii) VMB-AS release more EVs per cell than STR-AS under basal conditions; and (iii) only VMB-AS respond to CCL3 by producing more EVs, suggesting differential AS-EV secretion rate according to PD brain region. Next, addressing AS-EV potential against oxidative stress and mitochondrial toxicity, we found that AS-EVs, especially CCL3-AS-EVs, fully counteract H_2_O_2_-induced caspase-3 activation. Furthermore, using high resolution respirometry, we demonstrated that AS-EVs rescue the neuronal mitochondrial complex I function impaired by MPP^+^, with VMB-AS-EVs fully restoring ATP production in MPP^+^-injured neurons, highlighting a regional diversity of AS-EVs with neuroprotective implications for PD.

## 1. Introduction

EVs are nanometric (30-1000 nm) lipid membranous structures released by virtually all cell types into the extracellular milieu, where they can be captured by adjacent or distal cells.^[1–3]^ EVs is a general term used to describe a complex set of vesicles with distinct biogenesis and release mechanisms. Exosomes, microvesicles, and apoptotic bodies partially overlap in terms of dimension and composition, making it difficult to identify specific EV subclasses.^[4]^ Based on size, they are referred to as medium-large EVs (> 200 nm) or small EVs (< 200 nm).^[4,5]^ The importance of EVs in mediating cell-to-cell communication resides in the ability to deliver different cargoes (i.e., nucleic acids, proteins, metabolites, lipids) to target cells, thus influencing their fate.^[6–8]^ EVs have been identified in body fluids as potential new biomarkers for several diseases and also exploited as advanced nanotherapeutics in regenerative medicine.^[9,10,19,20,11–18]^ Like their synthetic liposomal counterpart, EVs protect their payloads from the action of nuclease and protease, allowing either a long distance delivery. Differently from synthetic nanoparticles, EVs display innate properties, such as the ability to cross biological barriers (e.g., blood brain barrier), and a low immunogenicity. Moreover, EVs possess a fingerprint, inherited from their donor cells, which distinguish EVs derived from different cell types. ^[13,21–23]^ Importantly, the EV content reflects the “status” of the donor cell and can change in response to specific modifications in the microenvironment.^[24]^

EVs have been demonstrated to play several roles in physio-pathological conditions.^[25]^ In the context of neurodegenerative diseases, including Parkinson’s disease (PD), EVs were initially identified as vehicles of misfolded proteins,^[26–28]^ but in line with the dual role played by glial cells, EVs have been demonstrated to play also important neuroprotective functions.^[29,30]^ Astrocyte (AS) dysfunction is increasingly emerging as a critical feature of PD,^[31–42]^ the second most common neurodegenerative disorder, with no cure available to stop or reverse its progression.^[43]^ PD is characterized by the selective and unrestrained death of dopaminergic (DAergic) cell bodies of the substantia nigra pars compacta (SNpc), residing in the ventral midbrain (VMB).^[43–45]^ As a consequence, in the striatum (STR), DAergic terminals slowly degenerate leading to the classical motor features of PD (i.e. bradykinesia, rest tremor, rigidity and postural instability).^[43–46]^ Along with the chronic, age-dependent nigrostriatal degeneration, the abnormal accumulation of intraneuronal inclusions enriched in aggregated α-synuclein (α-syn), known as Lewy bodies (LBs) and Lewy neurites (LNs), and a massive astrogliosis, represent the major histopathologic hallmarks of the disease.^[46–49]^ The causes and mechanisms of DAergic neuron death still remain elusive, albeit current evidence implicate a complex interplay between several genes and many environmental factors, especially ageing, inflammation and oxidative stress, all robustly impacting the astroglial cell compartment.^[31,32,35–37,39,50–54]^ Particularly, converging data point to mitochondrial dysfunction as the pivotal final pathway for PD neurodegeneration, closely related to the selective vulnerability of nigrostriatal neurons and to the specific properties of the astroglial microenvironment.^[54–65]^ In fact, AS are active mediators of either beneficial or detrimental functions during neuronal degeneration, via the expression of a plethora of proinflammatory/anti-inflammatory molecules and neurotoxic/neuroprotective mediators.^[32,35,36,62,63,66,67]^ The balance between these messengers, together with the bidirectional signaling with microglial cells, will determine the fate towards a reparative process or a neuronal failure.

Accordingly, within the VMB, AS exert potent neuroprotective effects onto the vulnerable SNpc-DAergic neurons [reviewed in ^[64]^]. In particular, reactive VMB-AS were identified as main actors linking neuroinflammation to DAergic neuroprotection and repair in the 1-methyl, 4-phenyl, 1,2,3,6 tetrahydropyridine (MPTP) mouse model of basal ganglia injury.^[68]^ In this context, a wide gene expression analysis identified a major upregulation of certain chemokines, in particular CCL3, as important players for DAergic neurogenesis, survival and immunomodulation.^[35,36,68–70]^ Notably, *in vitro* studies unveiled CCL3-activated AS-neuron crosstalk as a critical element promoting both neuroprotection and neurogenesis from adult neural stem cells (NSCs).^[35,64,68]^

However, the molecular details of this complex intercellular signaling are still a matter of intense debate. The secretion of AS-derived extracellular vesicles (AS-EVs) represent a likely way of communication for AS towards injured DAergic neurons, thus providing a powerful tool for neuroprotection.^[30]^

## 2. Results

### 2.1. Secretion of AS-small EVs is region-specific

To assess potential differences among the two principal brain regions affected in PD, primary AS cultures were established from the VMB and STR of P2-3 mice (Figure S1, Supporting information) as described in.^[68]^ First, AS culture preparations were characterized under both basal and after exposure to the chemokine CCL3. The purity of AS cultures was confirmed by immunofluorescence (IF) staining showing ≥ 98% of GFAP^+^ cells and less than 2% of IBA1^+^ microglia in all the conditions (Figure S1A, C, Supporting information). AS proliferation was addressed using triple GFAP/BrdU/DAPI staining, indicating no changes in the percentage of proliferating AS (GFAP^+^ over the total number of BrdU^+^ cells) between the two brain regions, both in the absence or the presence of CCL3 (Figure S1B, D, E, Supporting information). Finally, the expression of the receptors for CCL3 (i.e., Ccr1 and Ccr5) was confirmed using qPCR in both VMB- and STR-AS cultures, showing no differences between experimental groups (Figure S1F, Supporting information), thereby establishing the conditions for AS-EVs characterization.

EVs were isolated from AS supernatants by differential centrifugation ^[71,72]^ and analyzed through a combination of different techniques, in order to evaluate dimensions, secretion rates and specific markers. As a first line of characterization, we performed nanoparticle tracking analysis (NTA) on all EV samples (**Figure 1**). The data displayed an enriched population of vesicles with a peak size around 100 nm, which is in the size range of small EVs (VMB-AS-EVs: basal, 97.6 ± 4.5 nm; CCL3-treated, 95.4 ± 5.9 nm; STR-AS-EVs: basal, 101.9 ± 1.3 nm; CCL3-treated, 105.8 ± 4.9 nm (Figure 1A).

**Figure 1.**
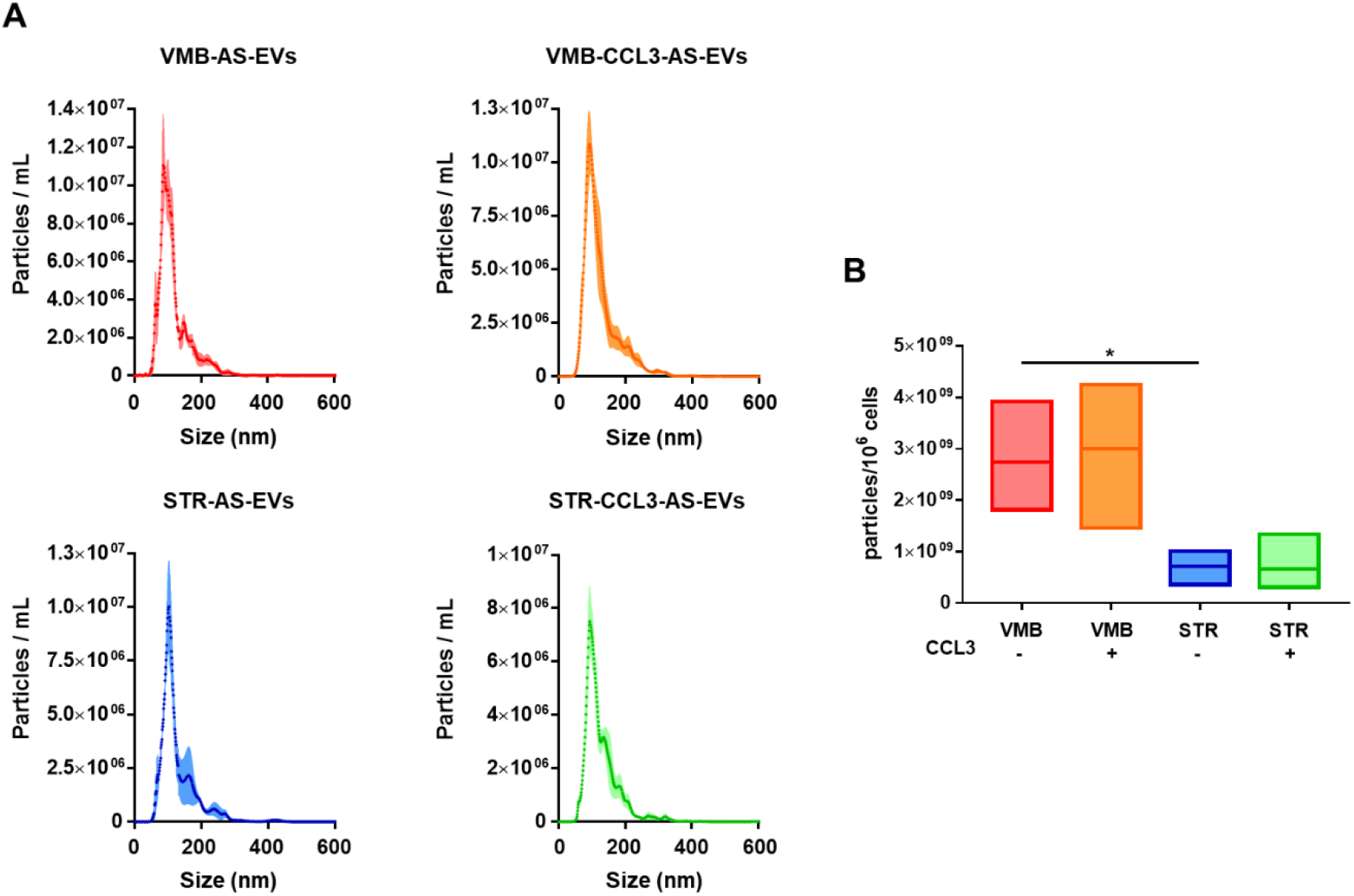
AS secrete vesicles enriched in the size range of small EVs. A) NTA analysis for size distribution (particles/ml) displays a peak around 100 nm. Error bars represent SEM from 3 independent replicates. B) Analysis of EV concentration (particles/106 cells) shows that in basal conditions VMB-AS secrete more EVs than STR-AS; VMB-CCL3-AS show a trend in releasing more EVs than basal. Data are presented as floating bars with line at mean from n=3 independent replicates. T-test *p < 0.05 (VMB-AS vs. STR-AS).

Although AS of all groups released vesicles with overlapping dimensions, we found a significant difference among AS from the two brain areas (Figure 1B), with VMB-AS releasing 3-4 fold more vesicles per million of cells. Moreover, CCL3 treatment induced an increased trend in EV release that was specific for VMB-AS (VMB-AS-EVs: 2.73×10^9^ ± 6.2 x10^8^; VMB-CCL3-AS-EVs: 3 x10^9^ ± 8.2 x10^8^; STR-AS-EVs: 7.13×10^8^ ± 1.9 x10^8^; STR-CCL3-AS-EVs: 6.65×10^8^ ± 3.3 x10^8^). Overall, these data demonstrate that AS retain EV secretion characteristics defined by their brain area origins.

### 2.2. CCL3 stimulates EV secretion specifically for VMB-derived AS

To further investigate the EV size distribution and possible differences in the secretion rate, dependent on the brain region and/or on the chemokine treatment, we performed a transmission electron microscopy (TEM) analysis on the very same EV samples. The images show the presence of many small vesicles with the typical cup shape (**Figure 2A**). First, we measured the diameter (nm) and the area (nm^2^) of each vesicle. The average diameter for each tested condition was ~70-80 nm, while the average area was ~6000 nm^2^, without any significant difference between groups (Figure 2B and S2, and Table S1, Supporting information). Next, we quantified the vesicles. Again, we found that VMB-AS released a significantly higher number of EVs than STR-AS (Figure 2C). Interestingly, the treatment with CCL3 stimulated VMB-AS to secrete more EVs, confirming the trend observed with the NTA (Figure 2C). Again, STR-AS-EVs did not show any significant difference, further supporting the idea that the EV secretion rates, and their responsiveness to microenvironmental cues, are specific for each brain area.

**Figure 2.**
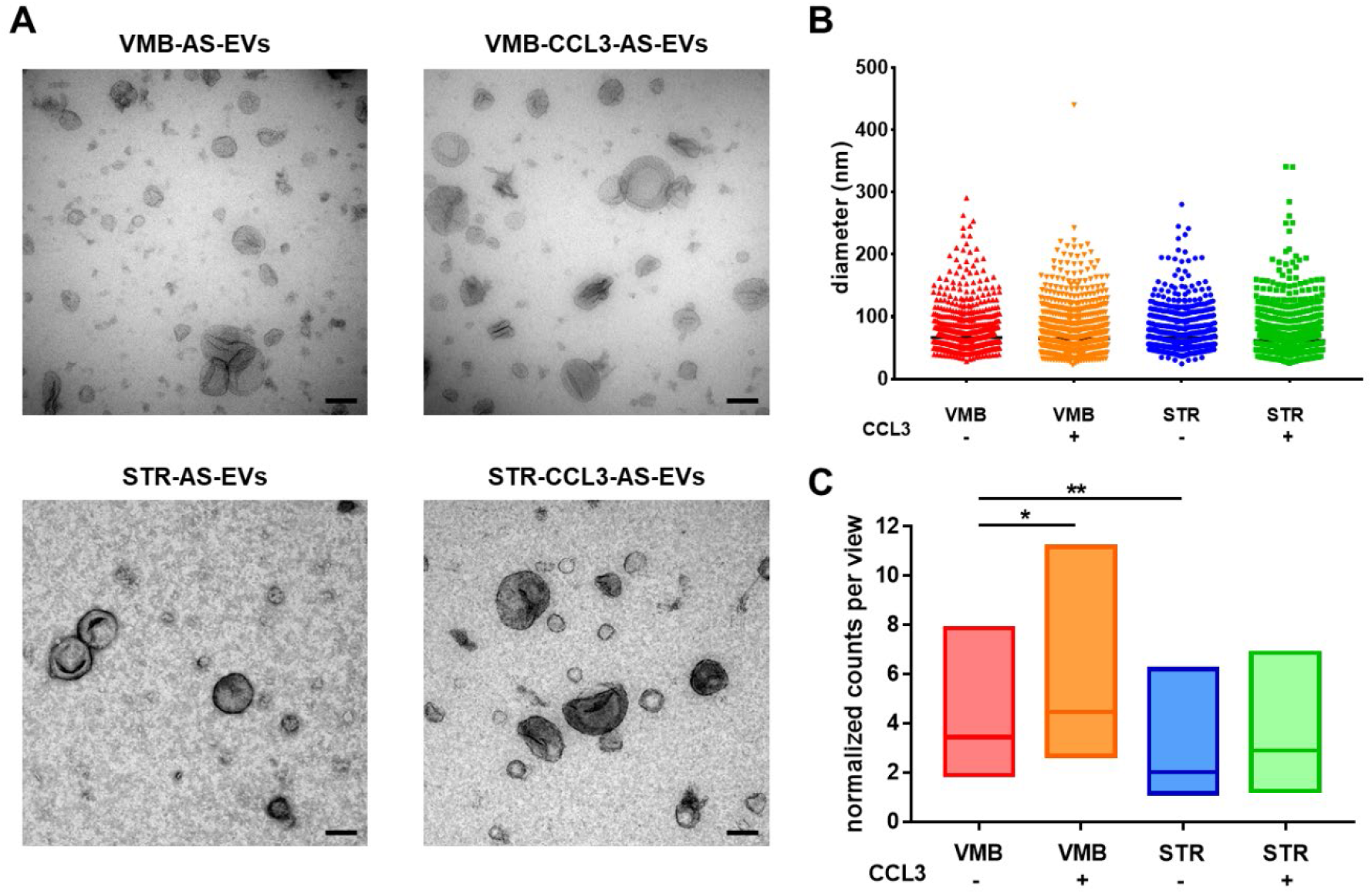
Brain area influences AS-EV secretion rate and responsiveness to CCL3 treatment. A) Ultrastructural analysis reveals the presence of small EVs secreted by AS in every condition. Scale bars: 100 nm. B) In all AS-EV samples the average diameter is around 70-80 nm. C) Quantitative analysis showed that in basal conditions VMB-AS secrete more EVs than STR-AS; the treatment with CCL3 stimulates VMB-AS to release more EVs, without any variation in terms of dimensions (VMB-AS-EVs: 4.27 ± 2; VMB-CCL3-AS-EVs: 5.68 ± 2.3; STR-AS-EVs: 2.5 ± 1.4; STR-CCL3-AS-EVs: 3.31 ± 1.8). Data are presented as scatter dot plots (B) or floating bars (C) with line at median from n=3 independent replicates. One-way ANOVA with Tukey’s multiple comparison *p < 0.05 (VMB-AS-EVs vs. VMB-CCL3-AS-EVs), **p < 0.01 (VMB-AS-EVs vs. STR-AS-EVs).

### 2.3. Both VMB- and STR-AS-derived vesicles are enriched in small EV markers

To further identify small EVs, we first applied an immunogold-labeling TEM approach for the small EV markers tetraspanins CD63 and CD9 on all AS-EV samples. The images in **Figure 3A-B** revealed the presence of both markers, visualized as well-defined 6 nm gold nanoparticles at the EV surface. In order to extend these results to other small EV markers, and exclude contamination of other cellular components, we used western blotting (WB) (Figure 3C-D). In line with the TEM data, we found a specific enrichment in the tetraspanins CD63/CD9 and Pdcd6ip (Alix) in all EV samples compared to donor AS. In contrast, the cellular markers Golga2 (for Golgi), Calnexin (for endoplasmic reticulum), SDHA (for mitochondria) and Actin (for cytoplasm), were mainly retained in the cells (Figure 3C-D). These results confirm that our vesicular preparations are enriched in small EVs.

**Figure 3.**
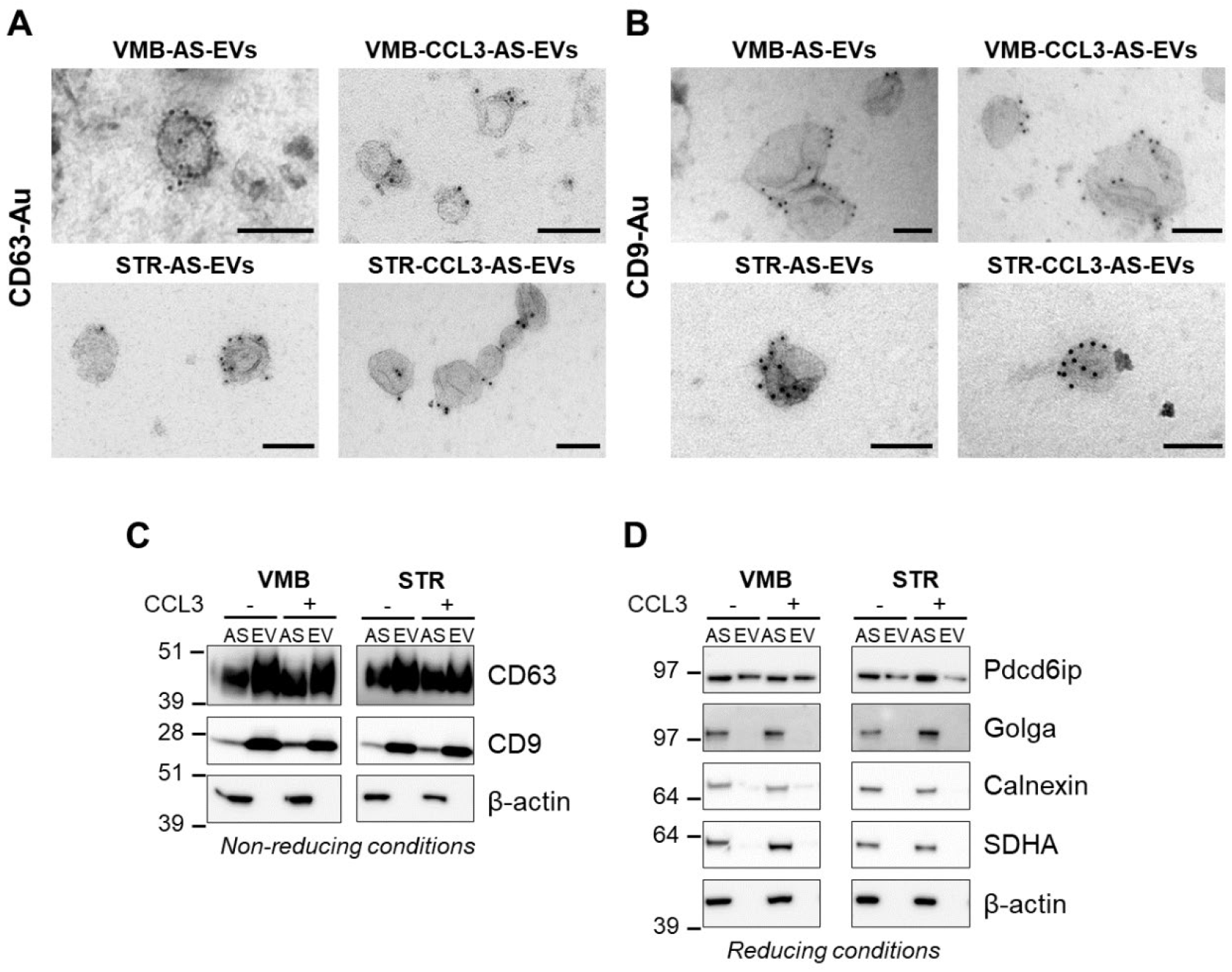
AS secrete vesicles enriched in small EV markers. A-B) Immunogold-TEM on EV samples with α-CD63 (A) and α-CD9 (B). Scale bars: 100 nm. C-D) WB analyses on EV lysates and corresponding AS donor cells. WBs for α-CD63/CD9 (C, in non-reducing conditions) and for Pdcd6ip (D, in reducing conditions) show an enrichment in the EV samples vs. donor AS. On the contrary, the cellular markers (i.e., Golga, Calnexin, SDHA and Actin) are mostly enriched in AS (D). All panels are representative of n=3 independent experiments showing the same trend.

### 2.4. CCL3-activated AS-EVs prevent H_2_O_2_ -induced caspase-3 activation in SH-SY5Y neurons

In order to verify whether AS-EVs display neuroprotective effects under neurodegenerative conditions, we selected the human neuroblastoma SH-SY5Y cell line as a model of neuronal target cells. The cells were differentiated to acquire a neuronal phenotype using 10 μM retinoic acid (RA) and gradual serum depletion, which stimulate neuroblastoma cells to extend neurites and to express tyrosine hydroxylase (TH).^[73]^ Firstly, we assessed the capacity of SH-SY5Y cell to internalize the AS-EVs (**Figure 4** and S3, Supporting information). We used two different approaches to label AS-EVs: (i) we labelled both VMB- and STR-AS with the PKH26 membrane dye, after which supernatants were collected and ultracentrifuged to isolate labelled EVs (Figure 4 and S3A, Supporting information); (ii) we used the PKH26 dye directly on AS-EVs after ultracentrifugation (Figure S3B, Supporting information). Both approaches were effective to obtain efficiently PKH26-labelled AS-EVs. Labelled EVs were then administered to target cells and their internalization was evaluated upon 6 h of incubation. The results show that AS-EVs are efficiently internalized by RA-differentiated SH-SY5Y cells (Figure 4 and S3, Supporting information). As further suggested by the orthogonal view analyses reported in Figure 4B, the PKH26-labelled AS-EVs are localized inside the cells and partially co-localize with TH (which has high affinity for phospholipid membranes).^[74]^ A volumetric 3D reconstruction of the AS-EVs intracellular distribution confirms the effective enrichment of AS-EVs within the cytoplasmic compartment (Figure S3A and Supplementary Movie 1-2, Supporting information). Moreover, the bright field/IF combined analyses suggest that AS-EVs are distributed in the whole cytoplasm, including neurite protrusions (Fig S3B, Supporting information). Overall, the internalization study supports that AS-EVs (and their cargoes) can be efficiently transferred to neuronal target cells (Figure 4 and S3, Supporting information).

**Figure 4.**
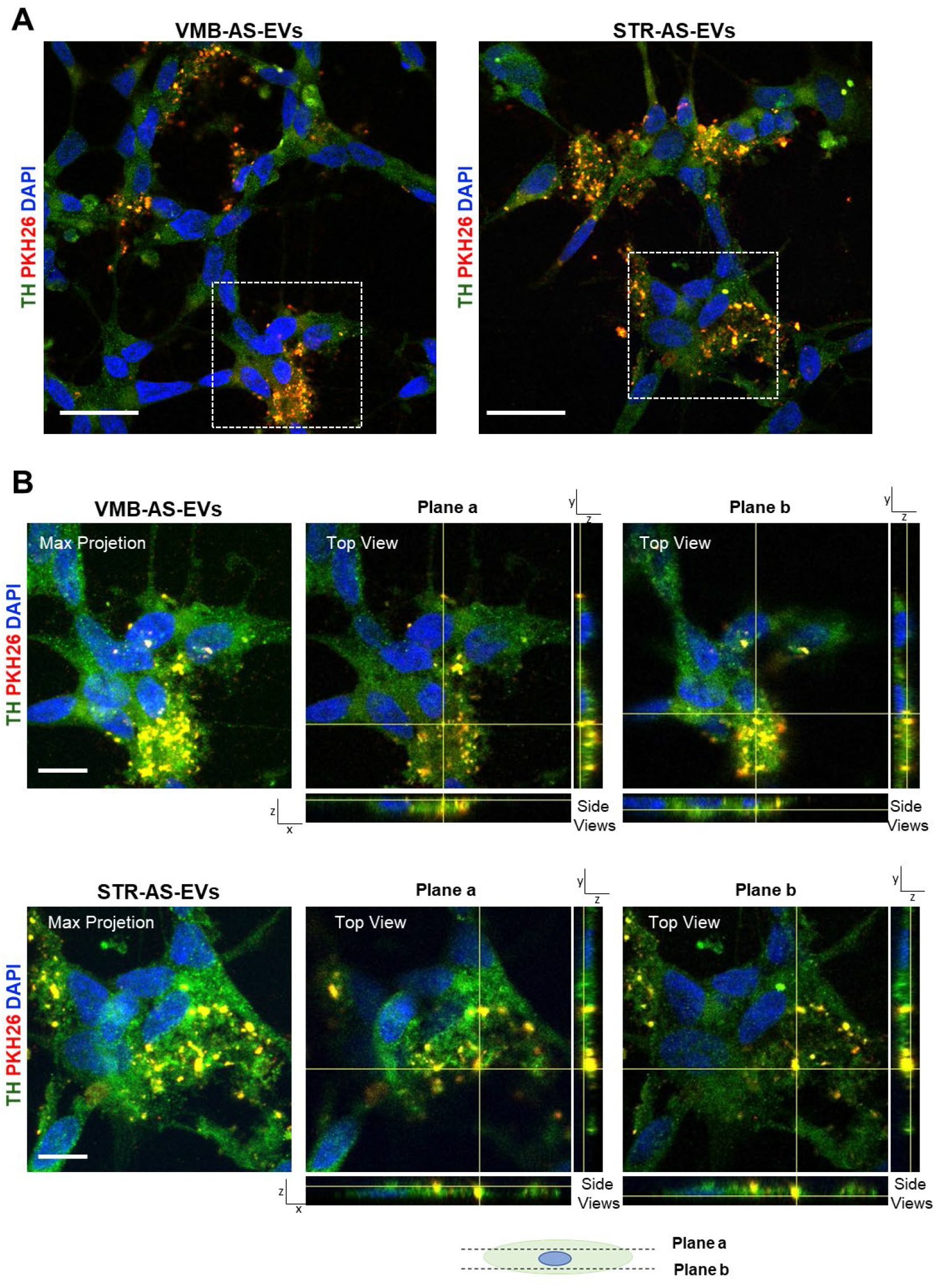
PKH26-labelled AS-EVs internalization within TH-positive RA-differentiated SH-SY5Y neuronal cells. A) Single plan confocal images show the uptake of both VMB-AS- and STR-AS-PKH26-labelled EVs by RA-differentiated SH-SY5Y. Scale bar 20 μm. B) Max projection and orthogonal views of representative fields (white dotted squares evidenced in (A). Each Max projection is composed of a stack of 15 individual z planes, acquired every 0.4 μm along the z axis. Scale bar 10 μm. Plane a and Plane b orthogonal views represent respectively two selected planes located above and below the cellular nuclei (along the z axis), as represented by the cellular schematic. In all panels PKH26 is in red, TH in green, whereas nuclear DAPI counterstain is in blue. Confocal images show that PKH26 labelled EVs are present within the cellular bodies of SH-SY5Y target cells.

To evaluate the effects of AS-EVs on target SH-SY5Y cells under oxidative stress conditions, we first used hydrogen peroxide (H_2_O_2_), a general source of ROS, and well recognized cytotoxic stimulus studied in different cellular models of PD. We selected the concentration of 35 μM, which reduces cell viability by ~40% (Figure S4, Supporting information) and focused on the key cell death executioner, cleaved caspase-3. We applied AS-EVs to the cells 6 h prior the challenge with H_2_O_2_ and the levels of cleaved caspase-3 were evaluated 24 h after H_2_O_2_ treatment via IF (**Figure 5**). Analysis of fluorescence intensity revealed that H_2_O_2_ induced a 3-fold increase of cleaved caspase-3 compared to untreated cells (CTRL), while the presence of both VM-B and STR-AS-EVs significantly reduced apoptosis levels (Figure 5A-B). Notably, only EVs from CCL3-treated AS fully rescued the apoptosis levels induced by H_2_O_2_ (Figure 5A-B).

**Figure 5.**
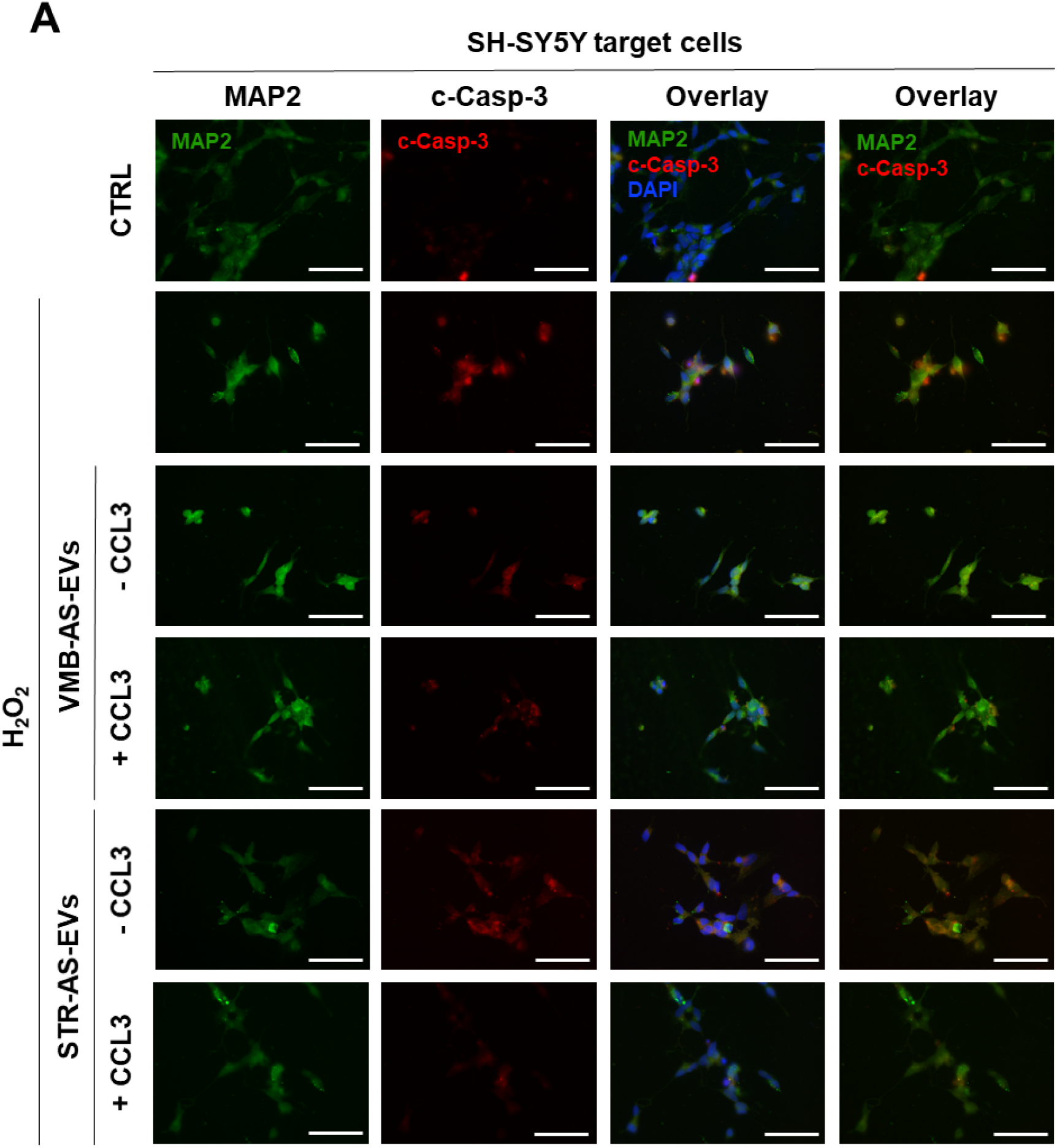
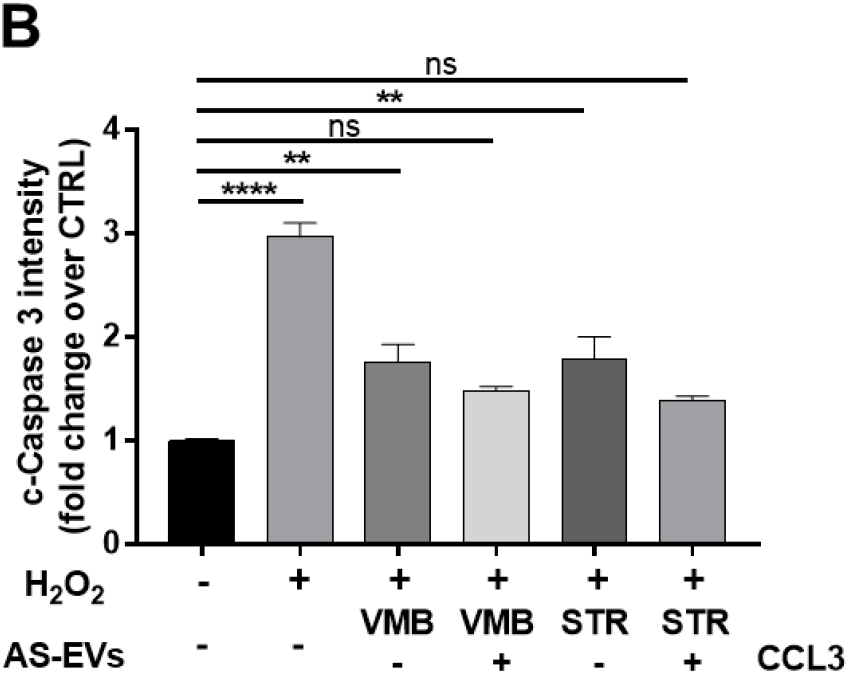
AS-EVs significantly reduce apoptosis in SH-SY5Y neurons challenged with H_2_O_2_. A) IF staining for MAP2 (in green), cleaved caspase-3 (in red) and DAPI (in blue), on differentiated SH-SY5Y exposed to AS-EVs and treated with 35μM H_2_O_2_. Scale bars: 50 μm. B) Quantification of cleaved caspase-3 IF staining. The fluorescent intensities of the signals were normalized over the cell number; values for CTRL were set to 1 for comparison: CTRL: 1; +H_2_O_2_: 2.97 ± 0.13; +VMB-AS-EVs + H_2_O_2_: 1.76 ± 0.16; +VMB-CCL3-AS-EVs + H_2_O_2_: 1.48 ± 0.04; +STR-AS-EVs + H_2_O_2_: 1.76 ± 0.24; +STR-CCL3-AS-EVs + H_2_O_2_: 1.35 ± 0.01. Data are expressed as mean ± SEM from n=3 independent replicates. One-way ANOVA with Dunnett’s multiple comparison **p < 0.01, ****p < 0.0001 vs CTRL, ns: not significant.

To verify that the observed neuroprotection was related to the specific effect of AS-EVs, we performed two control experiments (Figure S4, Supporting information). First, we found that the direct treatment with CCL3 on SH-SY5Y cells did not recover the viability of H_2_O_2_-injured neurons (Figure S4B, Supporting information). Second, the same protocol used to purify AS-EVs was applied to medium only, to exclude any potential effect of contaminant vesicles (cont-EVs). The treatment with cont-EVs did not change the levels of cleaved caspase-3 intensity induced by H_2_O_2_ (Figure S4C-D, Supporting information). Together, these results further identify AS-EVs as an effective mean to deliver protective cargoes to SH-SY5Y H_2_O_2_-injured neurons and support chemokine-activated AS, as neuroprotective players in nigrostriatal degeneration.^[68]^

### 2.5. Both VMB- and STR-AS-derived vesicles preserve the activity of mitochondrial complex I in SH-SY5Y neurons injured by the neurotoxin MPP^+^

Next, we extended the study of the neuroprotective potential of AS-EVs to the same target cells, exposed to the neurotoxin MPP^+^, a well-established *in vitro* model of PD. MPP^+^ affects DAergic neurons inducing a parkinsonian-like phenotype mainly inhibiting the activity of the NADH-ubiquinone oxidoreductase (complex I) of the electron transport chain ^[75,76]^. Furthermore, as we recently demonstrated, the toxin compromises the overall integrity of the inner mitochondrial membrane (IMM), affecting ATP production via a mechanism independent of complex I inhibition.^[73]^ In this context, we performed a dose-response curve for MPP^+^ and selected the dose of 1mM, which causes only a small (~10%) reduction of cell viability.^[73]^ Here, we used the same concentration of MPP^+^ to evaluate the neuroprotective effect of AS-EVs in the presence of MPP^+^-injured mitochondria, and to exclude non-specific mitochondrial deficits caused by a massive cell death process. EVs were applied to target cells 6 h before the challenge with the neurotoxin and mitochondrial functionality was analyzed by HRR after 24 h. The complete respiratory profile (i.e., the O_2_ consumption profile of cells upon addition of substrates or inhibitors) was obtained (see **Figure 6A** for a representative trace of the control cells alongside with the detailed protocol) and the main respiratory states of differently treated samples were analyzed.

**Figure 6.**
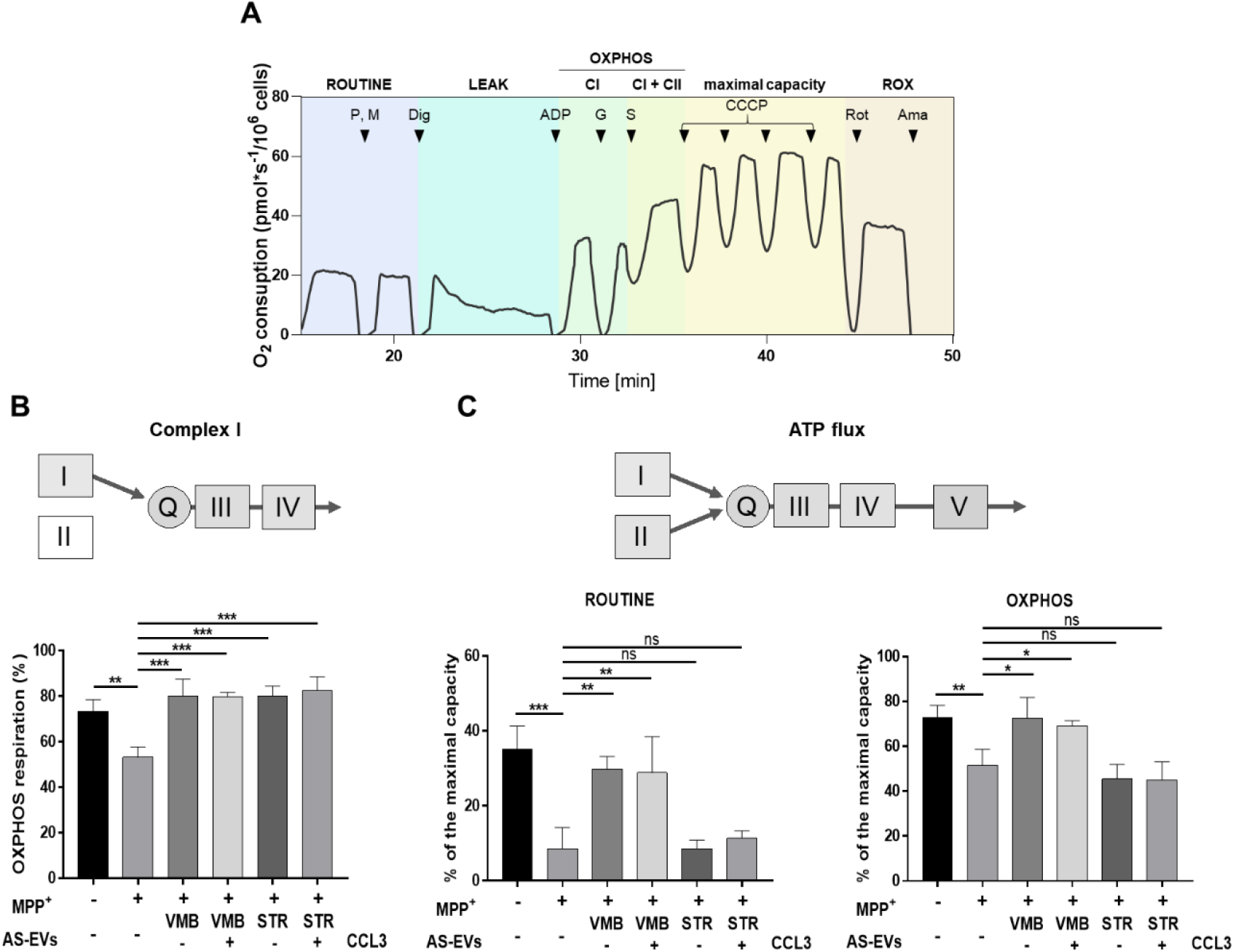
AS-EVs recover mitochondrial functions in SH-SY5Y neurons challenged with MPP^+^. A) Representative oxygraphic trace in untreated SH-SY5Y (control) cells alongside the specific protocol used. First, in intact cells, the physiological O_2_ consumption, corresponding to ROUTINE state, was measured. Second, adenylates were forced to leave the cells by a mild plasma membrane permeabilization in order to analyze the non-phosphorylating (or LEAK) state. Third, the contribution of complex I (CI) to the OXPHOS respiration was assayed in the presence of the sequential addition of the appropriate substrates (pyruvate, malate, glutamate) and a saturating ADP concentration. Then, addition of succinate allowed the activation of CII (CI + CII) and the achievement of total OXPHOS respiration. Fourth, the maximal capacity of the electron transport system (ETS) was obtained at the optimal uncoupler (CCCP) concentration. Fifth, the residual respiration (or ROX) was acquired after inhibition of ETS complexes with rotenone and antimycin a. P, pyruvate; M, malate; Dig, digitonin; G, glutamate; S, succinate; Rot, rotenone; Ama, antimycin a. B) Schematic representation of mitochondrial ETS complexes (upper panel) corresponding to the experimental set-up of the OXPHOS respiration measurement exclusively sustained by CI (lower panel). In this condition, e^-^flow through the ETS enzymes from CI but not from CII. The effect of MPP^+^ and/or EVs was tested in the same experimental conditions. The toxin reduces CI activity of about 30% compared to CTRL (CTRL: 73 ± 5 vs. MPP^+^ 53 ± 4, p=0.0014, n=4) while AS-EVs fully recover CI functionality of MPP^+^-treated SH-SY5Y cells (VMB-AS-EVs: 79 ± 7, VMB-CCL3-AS-EVs: 83 ± 7, STR-AS-EVs: 84 ± 11; 7, STR-CCL3-AS-EVs: 86 ± 8). Data are expressed as the ratio between OXPHOS driven by CI and total OXPHOS (driven by CI + CII) ± SD. One-way ANOVA with Tukey’s multiple comparison **p <0.01 (CTRL vs. MPP^+^), and ***p <0.001 (MPP^+^ vs. MPP^+^ + VMB-AS-EVs ± CCL3 and vs. MPP^+^ + STR-AS-EVs ± CCL3). C) Schematic representation of mitochondrial ETS and ATP synthase complexes (upper panel) corresponding to the experimental set-up of the ATP-related flux measurements in ROUTINE and OXPHOS state (lower panels). MPP^+^ reduces O_2_ flux devoted to ATP production compared to CTRL both in ROUTINE (CTRL: 35 ± 6 vs. MPP^+^ 10 ± 5) and OXPHOS states (CTRL: 72 ± 5 vs. MPP^+^ 51 ± 7). VMB AS-EVs promote a partial recovery of the flux in ROUTINE (VMB-AS-EVs: 30 ± 3 and VMB-CCL3-AS-EVs: 29 ± 10) and a complete recovery in OXPHOS (VMB-AS-EVs: 75 ± 9 and VMB-CCL3-AS-EVs: 69 ± 2). Data are expressed as percentage of the maximal ETS capacity ± SD. One-way ANOVA with Tukey’s multiple comparison. In ROUTINE: **p < 0.01 (MPP^+^ vs MPP^+^ + VMB-AS-EVs ± CCL3), ***p <0.001 (CTRL vs. MPP^+^). In OXPHOS: *p < 0.05 (MPP^+^ vs. MPP^+^ + VMB-AS-EVs ± CCL3), **p < 0.01 (CTRL vs. MPP^+^); ns: not significant.

Given the specific effect of MPP^+^ on complex I, we focused on its activity by analyzing the contribution of complex I to the overall oxidative phosphorylation (OXPHOS) respiration. This was achieved by two distinct steps: i) with the mild permeabilization of plasma membranes, allowing the exit of substrates and thus the complete inhibition of OXPHOS respiration; ii) with the stimulation of complex I activity with pyruvate, malate, glutamate and ADP at saturating concentration (Figure 6A). This set-up allows the electrons to flow from complex I – but not from complex II – to complex III, through the Q junction. Only the subsequent addition of succinate enables complex II to participate to the total OXPHOS respiration. As shown in Figure 6B, in CTRL cells complex I accounted for ~73% of the overall OXPHOS, while MPP^+^ reduced its activity up to a value of ~53% as expected. In this context, all AS-EV samples promoted a significant increase of complex I activity in MPP^+^-injured cells, up to fully restore its functioning as in CTRL cells (Figure 6B). Together, these data indicate the ability of AS-EVs from the most affected PD brain regions to efficiently and specifically preserve complex I activity, at a concentration range far below MPP^+^-induced massive cytotoxicity.

### 2.6. EVs secreted by VMB-AS ameliorate ATP production in MPP^+^-injured SH-SY5Y neurons

Given the potential of all AS-EV samples to protect complex I activity from MPP^+^, we further assessed whether they positively impact on other critical features of MPP^+^-induced mitochondrial dysfunction. For instance, the toxin does not affect the total O_2_ consumption recorded in the presence of endogenous substrates in intact cells (the so-called ROUTINE), neither the one observed in the presence of externally added substrates in permeabilized cells (the total OXPHOS), as we recently described.^[73]^ Accordingly, in our set-up, both ROUTINE and OXPHOS state were not specifically affected by the neurotoxin, and no significant changes were observed when MPP^+^ was used in combination with AS-EVs (Figure S5A-B, Supporting information). On the other hand, the neurotoxin treatment dramatically reduces the ATP-related fluxes, also known as net fluxes ^[73]^. With this perspective, HRR was used to analyze the effect of AS-EVs on the O_2_ flux exclusively devoted to ATP production. Results showed in Figure 6C confirm that MPP^+^ reduces both ROUTINE and OXPHOS net fluxes, by up to –75% and about –29% respectively (Figure 6C). However, treatment with EVs from VMB-AS – but not from STR-AS – significantly ameliorated the reduction in ADP phosphorylation of MPP^+^-injured SH-SY5Y cells, with no significant differences between untreated or CCL3-activated AS-EVs (Figure 6C). In particular, the use of VMB-AS-EVs promoted a partial and total recovery of the flux in ROUTINE and OXPHOS net flow, respectively (Figure 6C). Conversely, STR AS-EVs were not able to counteract MPP^+^ mitochondrial toxicity either in ROUTINE or in OXPHOS (Figure 6C). In line with these data, the degree of coupling between oxidative phosphorylation and electron flows (the coupling efficiency) was fully restored alongside with the increased ATP-related flows with VMB-AS-EVs only (Figure S5C-D, Supporting information).

Overall, these data indicate a regional specificity of VMB- vs. STR-derived AS-EVs in their ability to rescue the mitochondrial functional capacity of SH-SY5Y cells under MPP^+^ injury.

## 3. Discussion

Reactive astrocytes are increasingly emerging as key players in the parkinsonian brain, exerting both “beneficial” and “detrimental” effects.^[31–33,35–41,64,68,69,77,78]^ Especially, the heterogeneous nature of AS has been emphasized in earlier and more recent studies, showing regional AS differential responses to a panel of both genetic and environmental factors, including ageing, inflammatory or neurotoxin exposures, all crucial vulnerability conditions for PD.^[31,32,35,36,64]^ Yet, the modality of the intricate AS-neuron crosstalk still remains undefined. Among the multiple modes of intercellular communication, the secretion of EVs have emerged as a powerful tool for the exchange of information, capable of inducing multiple functional responses depending on the microenvironment.^[30,79,80]^ Notably, EVs are actively secreted by most cell types of the brain and have also been identified in body fluids working as potential new biomarkers for PD and other neurodegenerative diseases.^[9–14]^ They are also exploited more and more as advanced nanotherapeutics in regenerative medicine.^[9,13,15–20]^

Here, we show for the first time that this form of communication mediated by EVs exists for astrocytes derived from the nigrostriatal system. Moreover, we elucidate the different intrinsic capacities of astrocytes in terms of EV secretion and function, depending on the specific brain area of origin (i.e., ventral midbrain vs. striatum). Combining a set of high-sensitivity imaging and biochemical techniques, we show that AS from both brain areas release a population of vesicles enriched in the range of small-EVs, in line with the presence of small microvesicles and exosomes in our preparations.

Interestingly, in basal conditions, the EV secretion rate is specific for each brain area, with AS derived from VMB releasing more EVs per cell than the STR. These nigrostriatal-specific differences in AS-EV secretion have potential functional implications when vesicles are transferred to target cells. Importantly, we also demonstrated that the treatment with the chemokine CCL3, plays a critical role in the regulation of EV secretion. This AS activation strategy stems from a recent discovery by our research team, identifying a 6-fold upregulation of CCL3 in the VMB of PD mice, *in vivo*, during nigrostriatal degeneration and self-recovery, whereby reactive AS were defined as the key components of DAergic neurorescue pathways against MPTP/MPP^+^ injury.^[68]^ It seems, therefore, of interest that CCL3 affects EV secretion in a region-dependent fashion – with only the VMB-AS releasing more vesicles in response to the chemokine – in the absence of any influence dependent on cellular viability and/or proliferation differences. These data support the high level of AS heterogeneity in the CNS, whose regulatory mechanisms (e.g., transcriptional vs. epigenetic programs) remain outstanding open questions for the field.^[81–83]^

It follows that the molecular machinery able to orchestrate the distinct EV secretion rates – according to the brain region and the specific exposure to an inflammatory trigger – needs to be further elucidated. Also, these findings call for a deeper understanding of the functional implications of such a specific response to microenvironmental cues between VMB (where the DAergic neurons reside) vs. STR (where they project) for the pathogenesis of PD, and eventually for the development of new therapeutic avenues. In fact, while a plethora of studies has identified AS harmful factors, little is known on both the mechanisms driving the induction of pro-reparative states and their cellular/molecular effectors.^[84,85]^

Initially identified as possible neurodegeneration’s Trojan horse, AS-EVs recently emerged as important mediators of “beneficial messages” towards target neurons.^[86]^ For example, cortical AS were found to secrete neuroprotective EVs, which can be internalized by SH-SY5Y cells treated with paraquat, an herbicide implicated as a risk factor in PD, used as a neurotoxic oxidative stress.^[87]^ In a more recent study, AS obtained from whole brains were demonstrated to attenuate MPP^+^-induced cell death in SH-SY5Y cells and primary DAergic neurons.^[88]^

To investigate the functional impact of AS-EVs from the nigrostriatal system in degenerative conditions, we employed the differentiated human neuroblastoma SH-SY5Y cell line as an *in vitro* model of neuronal target cells. We exposed SH-SY5Y cells to two distinct sources of toxicity – H_2_O_2_ and MPP^+^ – mimicking oxidative stress and mitochondrial dysfunction found in PD. Intriguingly, we found that depending on the challenge used, AS-EVs differentially mediate neuroprotection on these target cells. In particular, both VMB- and STR-derived AS secrete EVs that are able *per se* to counteract the cell death induced by H_2_O_2_, a general source of ROS. In this regard, the vesicles obtained from AS treated with CCL3 showed the highest efficacy in preventing the activation of caspase-3 in SH-SY5Y cells. This novel finding sheds light on the mechanism of chemokine-mediated neuroprotection previously documented for AS derived from the nigrostriatal system ^[68]^ and further implicates a relevant role of the inflammatory microenvironment in amplifying the “beneficial” AS-EVs-mediated neuroprotection. Along these lines, *in vitro* studies showed that AS specific exposure to CCL3 – but not to TNF-α or IL-1β – successfully reverted aging-induced loss of AS neuroprotective properties against MPP^+^ cytotoxicity ^[34]^. In addition, the exposure of AS to CCL3 promoted neurogenesis and DA neurogenesis from adult midbrain neural stem cells (NSCs),^[68]^ and even reverted the aged-AS to a proregenerative state.^[89]^ On the contrary, the harmful microglial environment sharply inhibited subventricular zone NSCs, thus emphasizing the capacity of chemokine-activated AS in promoting DAergic neuron plasticity.^[35,36,68,70]^

Our data support the notion that the region-specific characteristics of AS are likely to play important roles in modulating DAergic neuron vulnerability, especially their response to oxidative stress and inflammation. Under neurodegenerative conditions, many adaptive changes occur within the astroglial cell compartment, aimed to increase the defense against oxidative stress and inflammation, to improve mitochondrial performance, to provide neurotrophic support, and/or to activate adult neurogenesis.^[35,36,62–64,67–69]^

The existence of such context-dependent AS properties further integrates with the known specific neuronal vulnerabilities shown to different neurotoxic insults. In fact, DA is a recognized source of oxidative stress for SNpc neurons in the VMB ^[59,61,65,90]^ and DAergic terminals in the STR, that actively degenerate proportionally to increased levels of DA oxidation.^[90–92]^ Accordingly, VMB-AS are endowed with a major anti-oxidant system, i.e., nuclear factor erythroid 2 -like 2 (NFE2L2/ Nrf2)-antioxidant responsive element (ARE) axis, sufficient to protect against nigrostriatal degeneration in both neurotoxin and genetic models of PD.^[42,64,69,93]^ On the other hand, a deficiency of AS-Nrf2-ARE signaling is observed with age, MPTP or α-synuclein expression and cooperates to impair neurogenesis ^[35,94,95]^ and to promote long-lasting nigrostriatal toxicity with no recovery.^[34,69,96]^

Considering the specificity of AS responses to different injuries, we next used the active metabolite of MPTP – a well recognized neurotoxin recapitulating Parkinsonian symptoms in humans, non-human primates, and mice ^[76,97–99]^ – to investigate the ability of AS-EVs to target mitochondrial function. After conversion of MPTP into its active metabolite, MPP^+^ is sequestered in mitochondria where it promotes a selective inhibition of complex I of the electron transport chain. Indeed, the well described deficiency of mitochondrial complex I activity in the SNpc of patients with sporadic PD accounts for the majority of neuron loss ^[59]^. We found that a preventive treatment with AS-EVs from both VMB and STR efficiently restored complex I activity in neuronal target cells, severely affected by the toxin treatment. On the other hand, only EVs released by VMB-AS fully preserved mitochondrial functionality, as demonstrated by the analysis of oxygen flows devoted to ATP synthesis in intact and permeabilized cells. In fact, MPP^+^ compromises the net respirations via a general reduction of the IMM integrity.^[73]^ This is critical feature for the maintenance of the proton gradient in the IMS, and therefore essential for the ADP phosphorylation process. In MPP^+^-injured neurons, a part of the gradient is dissipated, by-passing ATP synthases, and the positive effect exerted on complex I is either nullified or not culminated in ATP production, as in STR-AS-EVs treated cells. Different intriguing factors may contribute to this novel distinct effect of VMB vs. STR AS-EVs, which need to be further addressed. Again, this specificity may depend on the particular brain area facing region-specific neuronal vulnerabilities and/or specific tasks. For example, in the VMB, SNpc neurons are selectively vulnerable to mitochondrial complex I inhibitors, vs. the exquisite and “mysterious” sensitivity of STR cell bodies to succinate dehydrogenase (SDH, mitochondrial complex II) inhibitors, such as the plant-derived mitochondrial toxin, 3-nitropropionic acid, 3-NP, causing striatal damage reminiscent of Huntington’s disease (HD).^[100]^

Overall, it seems tempting to suggest the involvement of EVs in the astrocyte-neuron crosstalk, in a region-specific and context-dependent way. However, more work is required to clarify whether AS possess the intrinsic capacity to engage EV-mediated functional interactions with more relevant target cells – including primary DAergic neurons and NSC – and to what extent this type of intercellular communication contributes to AS neuroprotective effects highlighted in experimental PD models ^[101]^. More importantly, future studies will elucidate the nature of such AS-driven EVs cargoes and their possible link with the mechanisms regulating the selection/trafficking of specific effectors towards EVs. Indeed, a key aspect to be more deeply explored will involve the identification of the molecular cargoes of AS-EVs, capable to promote neuronal resilience and NSC reactivation. A very long list of molecules (DNA, RNAs, proteins, lipids, metabolites etc.) have been identified over the years within EVs, whose relative abundance (vs. donor cells) changes in response to specific stimuli.

Understanding the specific relationship between such a long list of potential EV-shuttled candidates and the key pathways in neural target cells remains a challenge for the field. Also, the way(s) used by AS-EVs to interact with target cells need to be further characterized, with different options available, such as signaling upon membrane-to-membrane interaction or after receptor-mediated endocytosis – and their eventual intracellular fate. This information will be very useful to fully understand the physiological function of AS-EVs, as well as help to better diagnose CNS diseases and identify their therapeutic potential.

## 4. Conclusion

This work highlights a novel role for AS-EVs in the propagation of specific intercellular signaling to neurons, in a region-specific and context-dependent fashion. Importantly, AS-EVs from the most affected brain regions in PD are capable to counteract neuronal cell death by targeting mitochondrial neuroprotection. Further elucidation of the molecular AS-EV signatures will represent a significant advance in the understanding of the multiple levels of interaction that are established between the cells in the brain. In the long term, tailored AS-EVs to prevent disease progression and promote neurological recovery may be foreseen, with implications for the etiopathology and the treatment of PD and other neurodegenerative diseases. In this regard, the innovative combination with synthetic lipid nanostructures, ^[13,102]^ may results in the production of vesicles with a better cargo loading, a sustained release of the desired cargo, while preserving the natural and innate properties of cell-derived EVs.

## 5. Experimental Section

### 5.1. Primary astrocyte cultures and treatments

Wild type C57BL/6 animals were purchased from Charles River (Italian Minister of Health authorization number 442/2020-PR). Primary astroglial cell cultures were prepared as described in ^[68]^. Briefly, AS were obtained from VMB and STR brain areas from mice at postnatal days P2-P4 and cultivated in DMEM (1 g/L glucose, Sigma Aldrich, D6046) supplemented with 10% FBS (Biowest, S1810), 2 mM L-glutamine (Sigma Aldrich, G7513), 2,5 μg/ml amphotericin B (Sigma Aldrich, A2942) and 1% penicillin/streptomycin (Sigma Aldrich, P0781) at 37°C and 5% CO_2_ for 13-17 days in 10 cm dishes specific for primary cultures (Corning, 353803). Loosely adherent microglial cells were then removed by shaking. Cells were washed with sterile PBS and allowed to grow for another two days or reseeded onto glass coverslips in 24-well plates for immunofluorescence (IF) analyses. Cells were washed and then treated or not with the CCL3 300 ng/ml (R&D, 450MA050), in DMEM medium supplemented with 10% FBS depleted of exosomes (System Biosciences, EXO-FBS-250A-1). Cells were maintained in this medium for 24 h before supernatant collection for EV purification. For IF analyses, AS were labelled with rabbit α-GFAP antibody (Dako, Z0334), while microglial cells were stained with goat α-Iba1 antibody (Novus, NB100-1028). AS proliferation was evaluated by 5-Bromo-2’-deoxyuridine (BrdU) incorporation assay. The day before fixation, BrdU 5 μM (Sigma Aldrich, 19-160) was added to cells. Proliferative cells were stained with mouse α-BrdU antibody (Sigma Aldrich, B8434). Donkey Alexa fluor secondary antibodies were used, and nuclei were stained with DAPI (Sigma Aldrich, 32670-5MG-F). IF images were acquired with Leica microscope and analyzed with Fiji Image J software 1.51n. RNA was isolated from AS using the miRNeasy Mini Kit (Qiagen, 217004). Total RNA quantity and purity were assessed with the NanoDrop ND-1000 instrument (Thermo Scientific) and cDNA synthesis was performed using the High-capacity cDNA reverse transcription kit (Applied Biosystem, 4368814). Gene expression was studied via qPCR with TaqMan™ Universal PCR Master Mix (Applied Biosystem, 4324018) and the following Taqman probes (Thermo fisher scientific): Ccr1: Mm00438260_s1; Ccr5: Mm01963251_s1. mRNA levels were normalized relative to Gapdh (Applied Biosystem, 4352339E). Samples were tested in triplicate on a QuantStudio 3 Real-Time PCR System (Applied Biosystem) and expressed as ΔCt.

### 5.2. EVs isolation and characterization

AS supernatants were collected and immediately centrifuged at 1000 x g at 4°C for 15 minutes in order to pull down residual cells/cell debris. Next, the supernatants were subjected to ultracentrifugation in a Sorvall WX100 (Thermo Scientific). The first ultracentrifugation was performed at 100,000 g at 4°C for 75 minutes, in ultra-cone polyclear centrifuge tubes (Seton, 7067), using the swing-out rotor SureSpin 630 (k-factor: 216, RPM: 23200). Then the pellet was washed with cold PBS and ultracentrifuged again at the same speed for 40 minutes in thick wall polycarbonate tubes (Seton, 2002), using the fixed-angle rotor T-8100 (k-factor: 106, RPM: 41000). The resulting pellets, containing AS-EVs, were resuspended in PBS (for NTA, electron microscopy and functional experiments) or in RIPA buffer (for WB characterization).

### 5.3. Nanoparticle tracking analysis (NTA)

AS-EVs were diluted in PBS and analyzed for particle size distribution and concentration on a Nanosight NS500 (Malvern Instruments Ltd, UK) fitted with an Electron Multiplication – Couple Device camera and a 532 nm laser. Sample concentration was adjusted to 10^8^-10^9^ particles /mL and measurements were performed in static mode (no flow) at an average temperature of 21±1°C. A total of 3 to 5 videos of 60 seconds were recorded for each independent replicate, loading fresh sample for each measurement. Videos were processed on NTA 3.2 and a detection threshold of 8 was used. The remaining settings were set to automatic. Total particle concentration for each EV isolate was determined by NTA and used to calculate the number of EVs released per cell.

### 5.4. EV negative staining for transmission electron microscopy

AS-EVs were fixed with 0.1% PFA in PBS for 30 min. 200 mesh formvar and carbon coated nickel grids were glow-discharged to make the surface grid hydrophilic. Fixed samples were placed on the grids for 7 min, samples were washed with ultrapure water and stained with 2% uranyl acetate for 7 min and examined at 80 kV on a FEI Tecnai G2 Spirit (FEI Company, Hillsboro, OR) transmission electron microscope equipped with a Morada CCD digital camera (Olympus, Tokyo, Japan). To measure the number of vesicles in EM, we counted 10 random 60 000 X fields, each from a different square from the 200-mesh grid from each condition ^[103]^. The results were normalized taking into account the following parameters: the number of starting cells, the resuspension volume after ultracentrifugation, the volume used in the microscope grid, and the area (μm^2^) of each field in the grid.

### 5.5. EV immunogold labeling for transmission electron microscopy

To increase the hydrophobic properties of the grids 200 mesh formvar and carbon coated nickel grids were glow-discharged. Grids were placed on a 10 μL drop of each sample for 7 min and washed with PBS. Nonspecific reactions were avoided using blocking solution containing normal goat serum for 30 min. Then, samples were washed in 0.1% BSAc (Aurion, Wageningen, the Netherlands) in PBS. Samples were incubated in 10 μL of 1:50 primary antibody (rat-anti-CD9 (BD, 553758); or rat-anti-CD63 (MBL, D263-3)) in 0.1% BSAc (Electron Microscopy Sciences) for 1 h. After, samples were washed in 0.1% BSAc and incubated in 1:50 goat-anti-rat 6nm gold particles (Abcam, ab105300) in 0.1% BSAc for 1 h in the dark. Grids were rinsed with 0.1% BSAc and fixed with 2% Glutaraldehyde for 5 min and washed with ultrapure water. Finally, negative staining with 0.4% uranyl acetate was performed for 7 min. Samples were examined at 80 kV on a FEI Tecnai G2 Spirit (FEI Company, Hillsboro, OR) transmission electron microscope equipped with a Morada CCD digital camera (Olympus, Tokyo, Japan).

### 5.6. Western blotting

AS and EVs extracts were processed as in ^[71,72]^. Briefly, AS and EVs were lysed in RIPA buffer and protein concentration was measured with *DC* Protein Assay (Biorad, 500-0116), using BSA as standard. The same amount of cell or EV lysates were then loaded into 4-12% Bis-Tris plus gels (Invitrogen, NW04125BOX) in reducing or non-reducing conditions. Afterwards, proteins were transferred onto PVDF membrane. All primary and secondary antibodies are listed in **Table 1**.

**Table 1.**
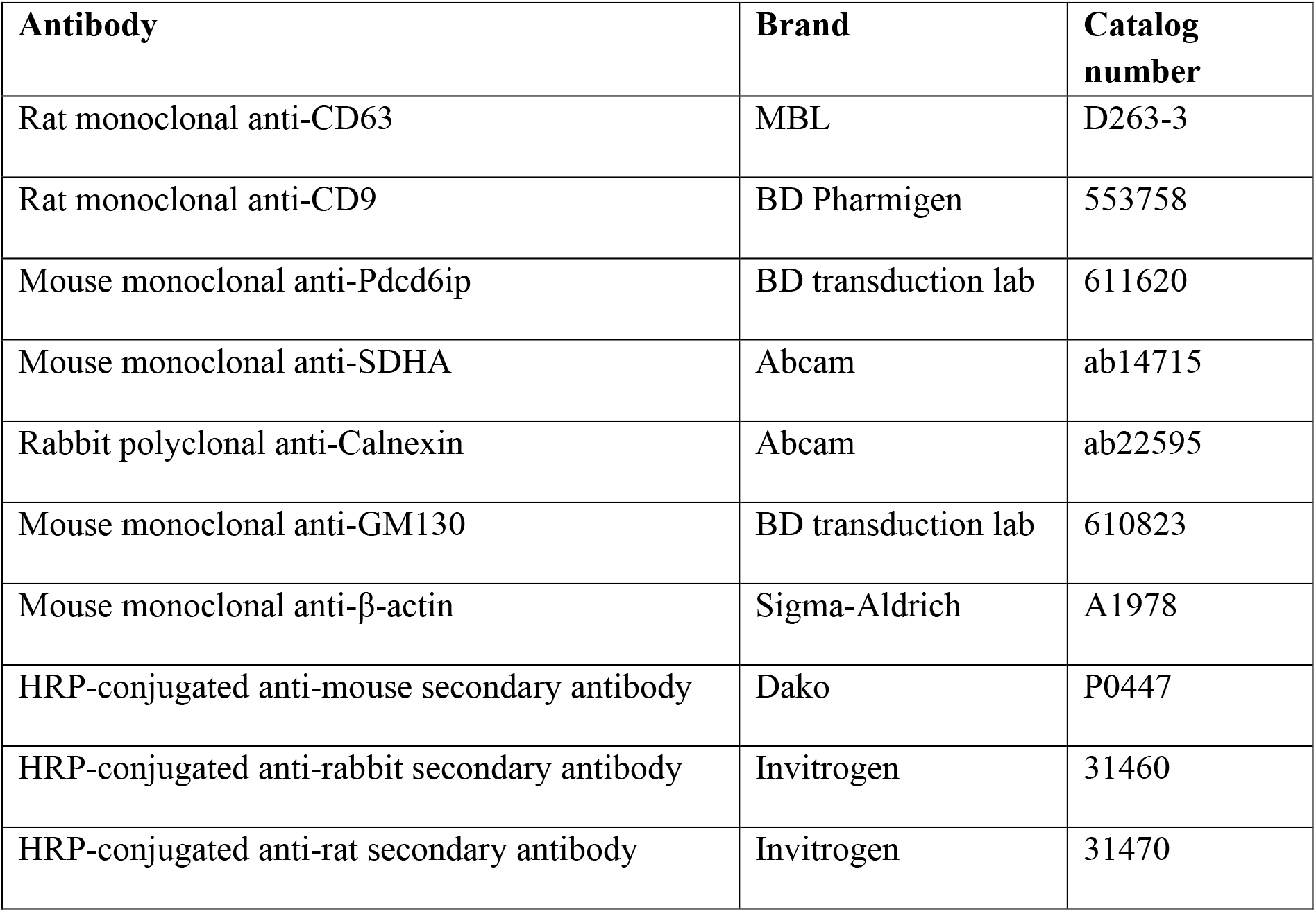
List of antibodies used in WB analyses.

| **Antibody** | **Brand** | **Catalog number** |
| --- | --- | --- |
| Rat monoclonal anti-CD63 | MBL | D263-3 |
| Rat monoclonal anti-CD9 | BD Pharmigen | 553758 |
| Mouse monoclonal anti-Pdcd6ip | BD transduction lab | 611620 |
| Mouse monoclonal anti-SDHA | Abcam | ab14715 |
| Rabbit polyclonal anti-Calnexin | Abcam | ab22595 |
| Mouse monoclonal anti-GM130 | BD transduction lab | 610823 |
| Mouse monoclonal anti-β-actin | Sigma-Aldrich | A1978 |
| HRP-conjugated anti-mouse secondary antibody | Dako | P0447 |
| HRP-conjugated anti-rabbit secondary antibody | Invitrogen | 31460 |
| HRP-conjugated anti-rat secondary antibody | Invitrogen | 31470 |

### 5.7. EV labeling

Prior to functional experiments, EV internalization was analyzed with two different approaches of EV labeling. First, AS were treated with the lipophilic dye PHK26 (Sigma Aldrich, MINI26-1KT), following the protocol suggested by the manufacturer. After 3 days cells were washed, and medium changed with DMEM supplemented with 10% FBS depleted of exosomes. EVs were isolated from AS supernatants after 24 h by ultracentrifugation. The resulting EVs were applied on the top of differentiated SH-SY5Y cells seeded onto polylysinated glass coverslips in 24 well plates. Target cells were stained with α-TH primary antibody (Millipore, AB152) as in.^[73]^ Imaging was performed using the confocal laser scanning microscope Leica TCS SP8. Image acquisitions were performed through LAS X software (Leica Microsystems). Image analyses were done using the open-source Java image processing program Fiji is Just ImageJ (Fiji). 3D reconstruction was done with the Fiji 3D Viewer dedicated plugin. For the second approach, EVs were directly labeled with the same lipophilic dye, following the protocol suggested by the manufacturer with some modification. Briefly, EVs derived from 90 ml of AS supernatant were ultracentrifuged, and the resulting pellets were resuspended in 0,3 ml of Diluent C with 4 μl of the red lipophilic dye and incubated at room temperature for 5 minutes, mixing every 30 second. The labeling was quenched by adding 1% BSA in PBS 1X and again ultracentrifuged. The resulting pellet, containing the labelled EVs, were resuspended in 100 ul PBS 1X. Residual PKH26 was eliminated into the Exosome Spin Column (thermo fisher scientific, 4484449) according to the manufacturer’s recommendations. Again, eluted EVs were applied on the top of differentiated SH-SY5Y cells seeded onto glass coverslips in 24 well plates. At the end of the treatment cells were fixed with 4% paraformaldehyde. IF images were acquired with Leica microscope and analyzed with Fiji Image J software 1.51n.

### 5.8. SH-SY5Y culture, differentiation, and treatments

SH-SY5Y cells were purchased from ICLC (Interlab Cell Line Collection, accession number ICLC HTL95013; obtained from depositor European Collection of Authenticated Cell Cultures (ECACC)) and cultured and differentiated as described in.^[73]^ Briefly, cells were maintained in MEM/F12 medium (Biochrom GmbH, F0325 and Sigma Aldrich, N4888) For cell differentiation, MEM/F12 was replaced with DMEM/F12 and 10 μM retinoic acid (Sigma Aldrich, R2625), and cultivated for 8 days with gradual serum deprivation until 0,5% FBS. At the end of differentiation, cells were detached and seeded at the density of 3 x 10^5^ cells/cm^2^ in 96-well (for dose response curve), 24-well (for caspase-3 IF staining) or 6-well (for HRR analysis) plates. Dose response curve for H_2_O_2_ was performed 24 h after the H_2_O_2_ treatment by using CellTiter Blue (Promega, G8080). CellTiter blue reagent (diluted 1:4 with PBS) was added directly to each well of 96 well plates and the plates incubated at 37°C for 4h. Then the fluorescent signal was read in Varioskan flash plate reader (Thermo fisher). For IF, cells were seeded on poly-L-lysine (Sigma Aldrich, P9155) coated glass coverslips. After two days, EVs were applied on the top of the cells by using the ratio 5:1 (i.e., EVs derived from five AS used to treat one SH-SY5Y cell). As a control, the vesicles eventually present as contaminants in the medium used to culture AS (cont-EVs) were also tested following the same passages used for AS-EVs. Briefly, AS culture medium was incubated for 24 h at 37°C and 5% CO_2_. Following ultracentrifugation, cont-EVs were resuspended in PBS and used to treat SH-SY5Y maintaining the same ratio with the starting volume of medium as for the purification of AS-EVs. 6 h later, cells were treated with H_2_O_2_ 35 μM for further 24 h. Coverslips were fixed with 4% PFA and stained with rabbit polyclonal anti-cleaved caspase-3 (Cell Signaling, 9664) primary antibody and with mouse monoclonal anti-map2 primary antibody (Merck Millipore, MAB3418). The secondary antibodies used were the anti-Rabbit Alexa Fluor 546 (Thermo fisher Scientific, A10040), and the anti-Mouse Alexa fluor 488 secondary antibodies (Thermo fisher Scientific, R37114). Nuclei were counterstained with DAPI. IF images were acquired with Leica microscope and analyzed with Fiji Image J software. The intensity of the cleaved caspase-3 signal was measured by using the following steps in ImageJ software: (i) analyze; (ii) measure; and (iii) integrated density, as in ^[104]^. Integrated density was normalized for the number of DAPI^+^ nuclei. All data are normalized over untreated control cells. As a further control, the chemokine CCL3 (at 30 ng/ml and 300 ng/ml) was added directly to SH-SY5Y cell cultures on 96-well plate 6 h before H_2_O_2_ exposure. Cell viability was evaluated 24 h after the H_2_O_2_ treatment with CellTiter Blue.

### 5.9. High-Resolution Respirometry (HRR) measurement

Capacity of different respiratory states in differentiated SH-SY5Y cells was assayed by High-Resolution Respirometry (HRR) using the O2k-FluoRespirometer (Oroboros Instruments). Cells were seeded in 6-well plates and, after two days, AS-EVs were applied on the top of differentiated SH-SY5Y cells by using the ratio 5:1, as before. 6 h later, cells were treated with MPP^+^ 1 mM for further 24 h. All the experiments were performed in mitochondrial respiration buffer Mir05 (Oroboros Instrument, 60101-01) at 37° C under constant stirring (750 rpm). A specific Substrate-Uncoupler-Inhibitor Titration (SUIT) protocol was used for the determination of the oxygen consumption in each specific respiratory state, as detailed in ^[73]^. Briefly, respiration in the presence of endogenous substrates or ROUTINE was measured in intact cells. The mild-detergent digitonin (Sigma Aldrich, D5628) was added at the final concentration of 4 μM in order to obtain the permeabilization of plasma membrane without compromising mitochondrial membranes integrity. The oxygen consumption after permeabilization or LEAK was determined in the presence of 5 mM pyruvate (Sigma Aldrich, P2256) and 2 mM malate (Sigma Aldrich, M1000) but not adenylates. The contribution of complex I to the OXPHOS respiration was achieved by addition of 10 mM glutamate (Sigma Aldrich, G1626) in presence of a saturating concentration of ADP (2.5 mM, Sigma Aldrich, 117105). The OXPHOS respiration was then stimulated with the addition of 10 mM succinate (Sigma Aldrich, S2378). The uncoupled maximal capacity of the electron transport system (ETS) was obtained after titration with 0.5 μM of uncoupler carbonyl cyanide 3-chlorophenylhydrazone (CCCP, Sigma Aldrich, C2759) up to the complete dissipation of the proton gradient. Finally, the residual oxygen consumption or ROX was obtained upon addition of 2 μM rotenone (Sigma Aldrich, R8875) and 2.5 μM antimycin A (Sigma Aldrich, A8674). The oxygen consumption in ROUTINE, LEAK, OXPHOS, and ETS capacity was corrected for the ROX. Values were then expressed as Flux Control Ratio (FCR) of the maximal respiration, using ETS capacity as a reference state ^[105]^. The oxygen flux related to ATP synthesis was determined by correcting ROUTINE and OXPHOS for the LEAK respiration. Coupling efficiencies were calculated by correcting each state for LEAK respiration and expressing it as a percentage of the capacity in that specific state.^[105]^

### 5.10. Statistical Analysis

GraphPad Prism software was used for all analyses. For IF on AS, images from n=4 independent biological replicates (from 4 to 10 images for each replicate) were analyzed by one-way ANOVA and Tukey’s multiple comparisons test, and expressed as mean (± SEM). For qPCR, data from n=3 independent biological replicates were analyzed by one-way ANOVA and Dunnett’s multiple comparisons test, and expressed as mean (± SD). For NTA, data from n=3 independent biological replicates (a total of 3 to 5 videos of 60 seconds recorded for each biological replicate) were analyzed by t-test, and expressed as mean (± SEM). For EM, data from n=3 independent biological replicates (10 fields for each replicate) were analyzed by one-way ANOVA and Tukey’s multiple comparisons test and expressed as mean (± SD). For dose-response curve of H_2_O_2_, data from n=3 independent biological replicates were analyzed by nonlinear regression, dose-response- inhibition ([Inhibitor] vs. response - Variable slope (four parameters)), and expressed as mean (± SEM). For IF on SH-SY5Y, data from n=3 independent biological replicates (two technical replicates, 8 images in total for each biological replicate) were analyzed by one-way ANOVA and Dunnett’s multiple comparisons test, and expressed as mean (± SEM). For HRR measurement, the following independent biological replicates have been performed: n=4 for CTRL and MPP^+^, n=3 for +/− VMB-AS-EVs, n=2 for +/− STR-AS-EVs. Data were analyzed by ANOVA and Tukey’s multiple comparisons test, and expressed as mean (± SD).

## Supporting Information

Supporting Information is available from the Wiley Online Library or from the author.

## Acknowledgements

The authors thank Aviva M. Tolkovsky, Vito De Pinto and Massimo Libra for critically discussing the article. We acknowledge the support and technical assistance of Nunzio Vicario. We are grateful to CAPiR (Center for Advanced Preclinical *in vivo* Research) Team for maintenance and care of animals. The authors acknowledge the PON project Bio-nanotech Research and Innovation Tower (BRIT), financed by the Italian Ministry for Education, University and Research (MIUR) (Grant no. PONa3_00136). We acknowledge the technical assistance of the BRIT Team with essential instruments. We are grateful to the Pharmacology and the Biology and Genetics sections at the BIOMETEC Department which host our laboratories.

The project has been supported by the “Brain to South” grant (Fondazione con il Sud – Bando Capitale Umano ad Alta Qualificazione 2015). The research program also received support from the Italian Ministry of Health (Cur. Res. And Finalized Res projects 2010-2020), from University of Catania (“Bando-Chance”, PRIN-2015, PIACERI and PhD program in Biotechnology). We acknowledge “AIM Linea 1 Salute (AIM1833071) to A. Magrì. We acknowledge the Nano-scaffolding for neuronal migration and generation project (PCI2018-093062) granted by the Spanish Ministry of Science, Innovation and Universities and Red de Terapia Celular (TerCel-RD16/0011/0026).

## Author contribution

L.L., F.L’E., A.Ma., N.F. L.P-J., S.P., J.M.G-V, A.Me, B.M. and N.I. conceived and designed the experiments. L.L., F.L’E., G.P., N.T., F.P. and N.I. performed primary astrocyte and SH-SY5Y cultures and treatments, EV isolation, EV labeling. L.L., F.L’E., G.P., S.V. C.T., S.C. and N.I. carried out immunofluorescence, confocal and western blotting analyses. L.L., C.A.P.B. and N.I. performed nanoparticle tracking analysis under the supervision of N.F. L.L. and M.J.U-N. performed electron microscopy analyses under the supervision of J.M.G-V. L.L., A.Ma, G.P. and P.R. performed high resolution respirometry analysis under the supervision of A.Me. L.L., A.Ma., M.J.U-N., G.P., S.V., C.A.P.B. and N.I. performed statistical analyses. L.L., A.Ma., B.M. and N.I. wrote the original draft. All authors contributed to the preparation of the figures and to the final version of the manuscript.

