## Supplementary materials for "Small Extracellular Vesicles Secreted by Region-specific Astrocytes Ameliorate the Mitochondrial Function in a Cellular Model of Parkinson’s Disease"

### Supporting Information

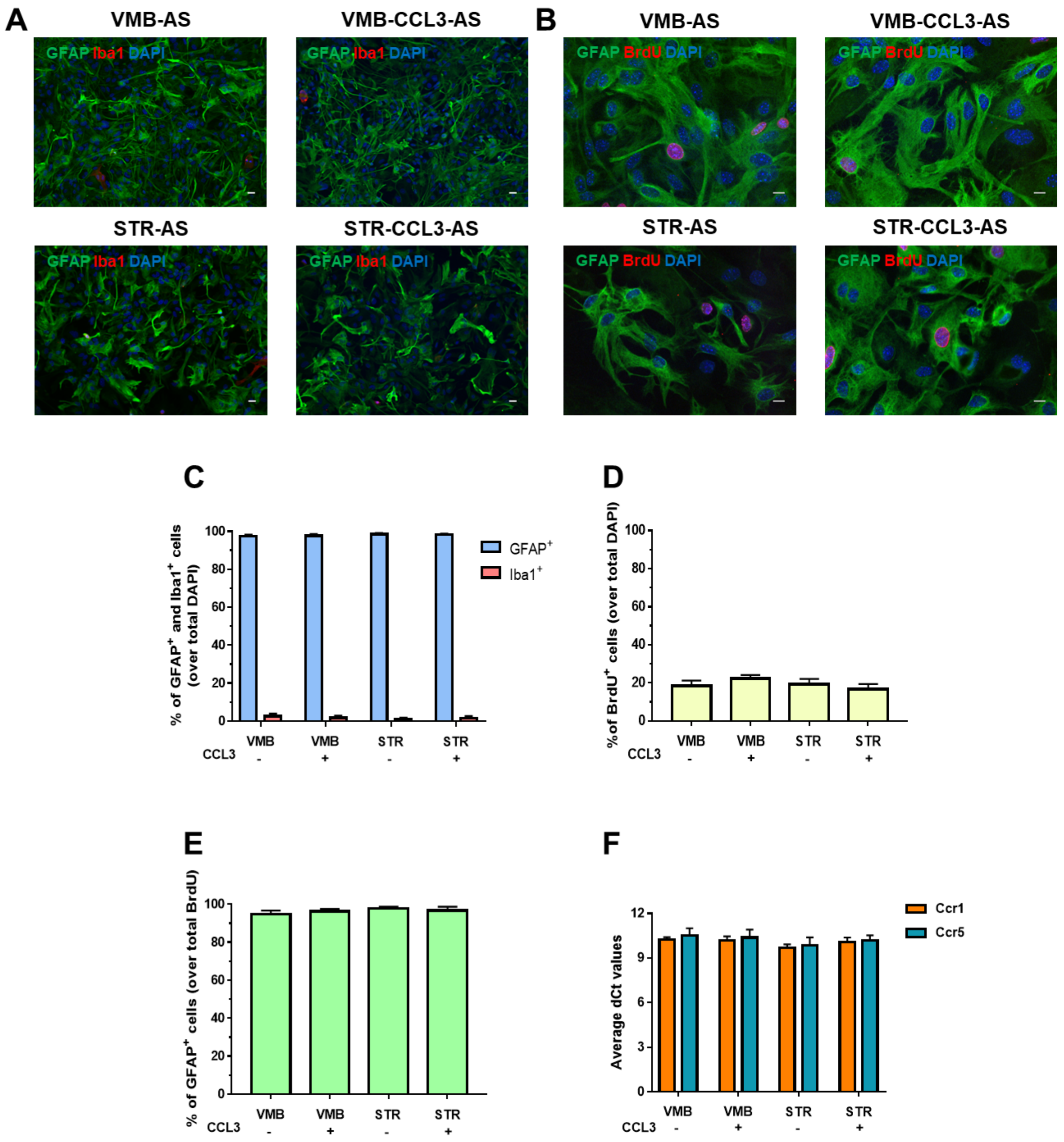


**Figure S1** Characterization of AS primary culture under basal and CCL3-treated conditions. A) IF images show the presence of AS (GFAP$^{+}$ cells, in green) and microglial cells (Iba1$^{+}$ cells, in red), over total DAPI$^{+}$ nuclei (in blue). Scale bars: 20 µm. B) IF images show the presence of proliferative AS (GFAP$^{+}$/BrdU$^{+}$ cells, in green and red respectively), over total DAPI$^{+}$ nuclei (in blue). Scale bars: 10 µm. C) Quantification of GFAP$^{+}$ cells over total DAPI$^{+}$ nuclei. d-e Quantification of BrdU$^{+}$ cells over total DAPI$^{+}$ nuclei (in D), or GFAP$^{+}$ cells over total BrdU$^{+}$ (in E). F) qPCR analyses of CCL3 receptors in AS, showing average dCt values for Ccr1 and Ccr5. Gapdh was used as housekeeping gene. Differences between experimental groups in c-f are not statistically significant. Data are presented as mean ± SEM from n=4 for IF, and n=3 for qPCR, independent replicates

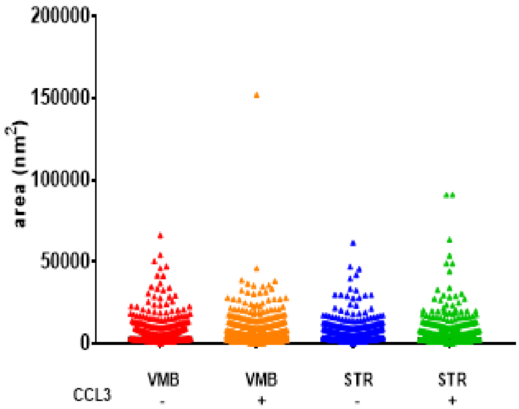


**Figure S2** All AS-EVs samples have an average area around 6000 nm$^2$. Data are represented as scatter dot plots with line at median from n=3 independent replicates

**Table S2** Diameter and area values of AS-EV samples.

| **Diameter (nm)** | **VMB-AS-EVs** | **VMB-CCL3-AS-EVs** | **STR-AS-EVs** | **STR-CCL3-AS-EVs** |
|---|---|---|---|---|
| Minimum | 28,28 | 22,12 | 24,19 | 25,37 |
| Maximum | 290,8 | 440,1 | 280,3 | 340,9 |
| Median | 66,58 | 64,65 | 66,37 | 61,83 |
| Mean | 78,9 | 76,38 | 79,05 | 73,77 |
| Std. Deviation | 38,62 | 39,57 | 37,74 | 39,75 |
| **Area (nm$^2$)** | **VMB-AS-EVs** | **VMB-CCL3-AS-EVs** | **STR-AS-EVs** | **STR-CCL3-AS-EVs** |
| Minimum | 628 | 384,3 | 459,4 | 505,6 |
| Maximum | 66429 | 152155 | 61689 | 91281 |
| Median | 3482 | 3283 | 3460 | 3002 |
| Mean | 6059 | 5810 | 6025 | 5513 |
| Std. Deviation | 7420 | 8150 | 6874 | 8192 |

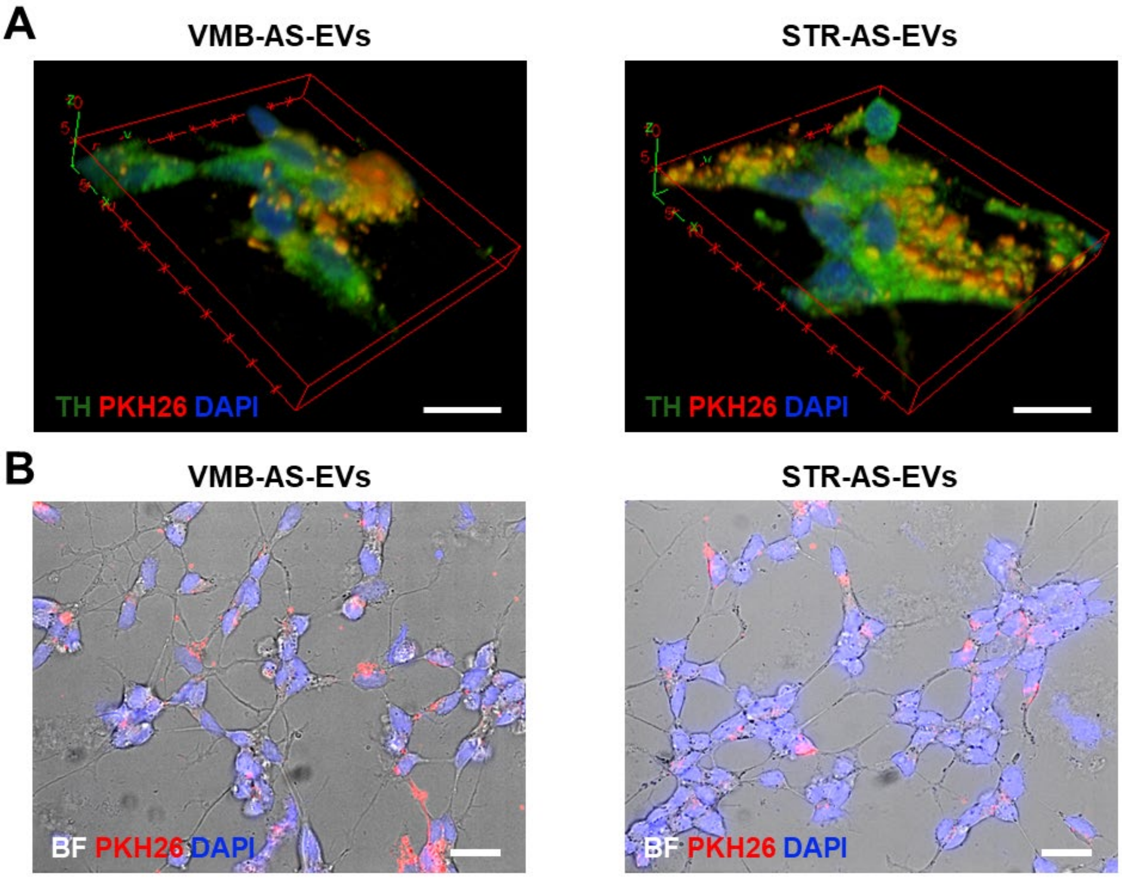


**Figure S3** Internalization of PKH26 fluorescently labelled EVs in RA-differentiated SH-SY5Y. A) 3D reconstruction from all z stacks (see Fig. 4B). Scale bars 10 µm. B) IF (in red PKH26 labelled AS-EVs and in blue DAPI counterstained nuclei) and bright field (whole cells) images of RA-differentiated SH-SY5Y upon treatment with PKH26-labeled EVs. EVs are distributed in cell bodies and also in neurites. Scale bars: 20 µM

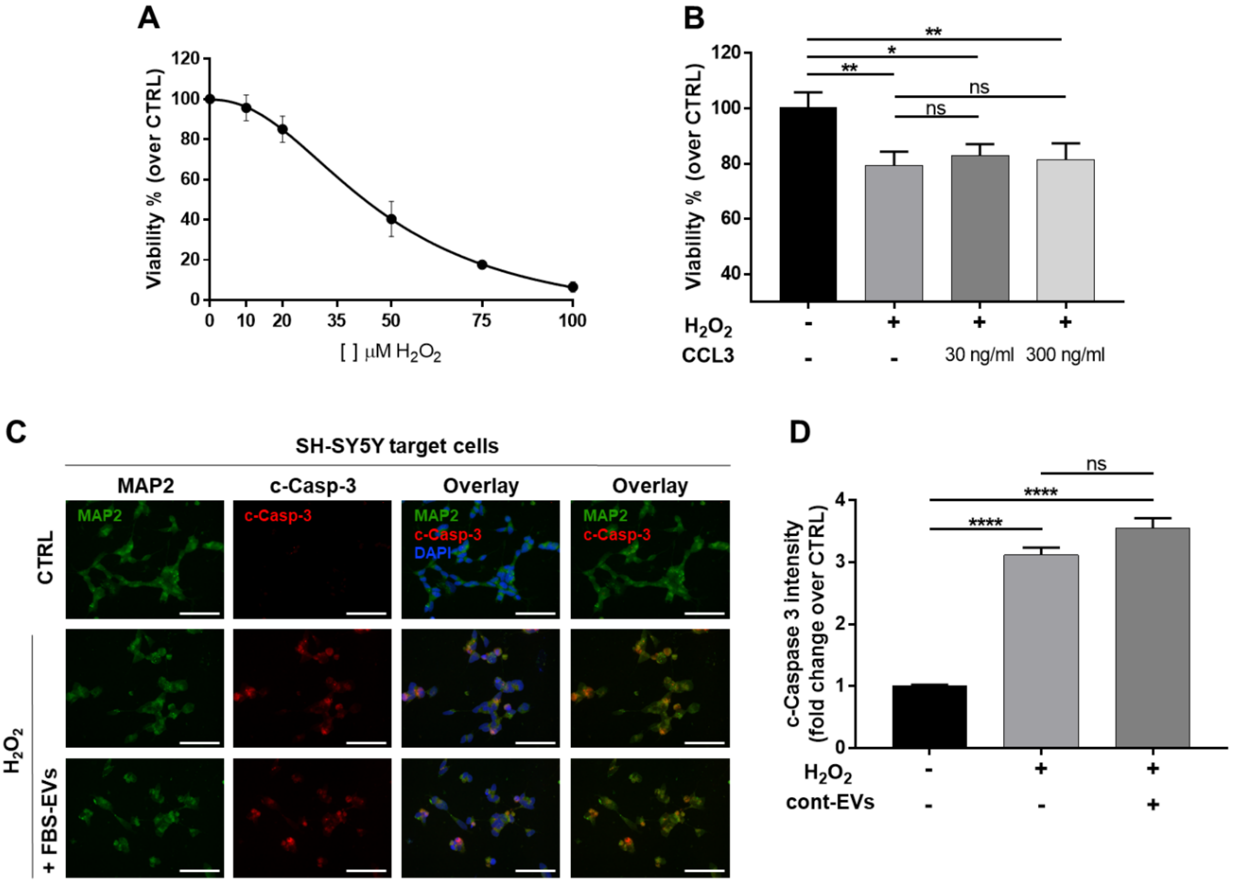


**Figure S4** Dose response curve of $H_2O_2$ and effects of CCL3 and FBS-EVs on $H_2O_2$-injured cells. A) Dose response curve of $H_2O_2$ on differentiated SH-SY5Y cells. B) Analysis of cell viability (% over CTRL) of CCL3-treated SH-SY5Y exposed to 35µM $H_2O_2$. Both concentrations of CCL3 do not affect viability of injured neurons. C) IF staining for MAP2 (in green), cleaved caspase-3 (in red) and DAPI (in blue), on differentiated SH-SY5Y exposed to cont-EVs and treated with 35µM $H_2O_2$. Scale bars: 50 µm. D) Quantification of cleaved caspase-3 IF staining. The fluorescent intensities of the signals were normalized over the cell number; values for CTRL were set to 1 for comparison. Data are expressed as mean ± SEM (in A, D) and SD (in B). One-way ANOVA with Tukey's multiple comparison. In (B) $*p < 0.05$ (CTRL vs. $H_2O_2$ + CCL3 30 ng/ml), $**p < 0.01$ (CTRL vs. $H_2O_2$ and vs. $H_2O_2$ + CCL3 300 ng/ml), ns: not significant. In (D) $****p < 0.0001$ (CTRL vs. $H_2O_2$ and $H_2O_2$ + FBS-EVs), ns: not significant

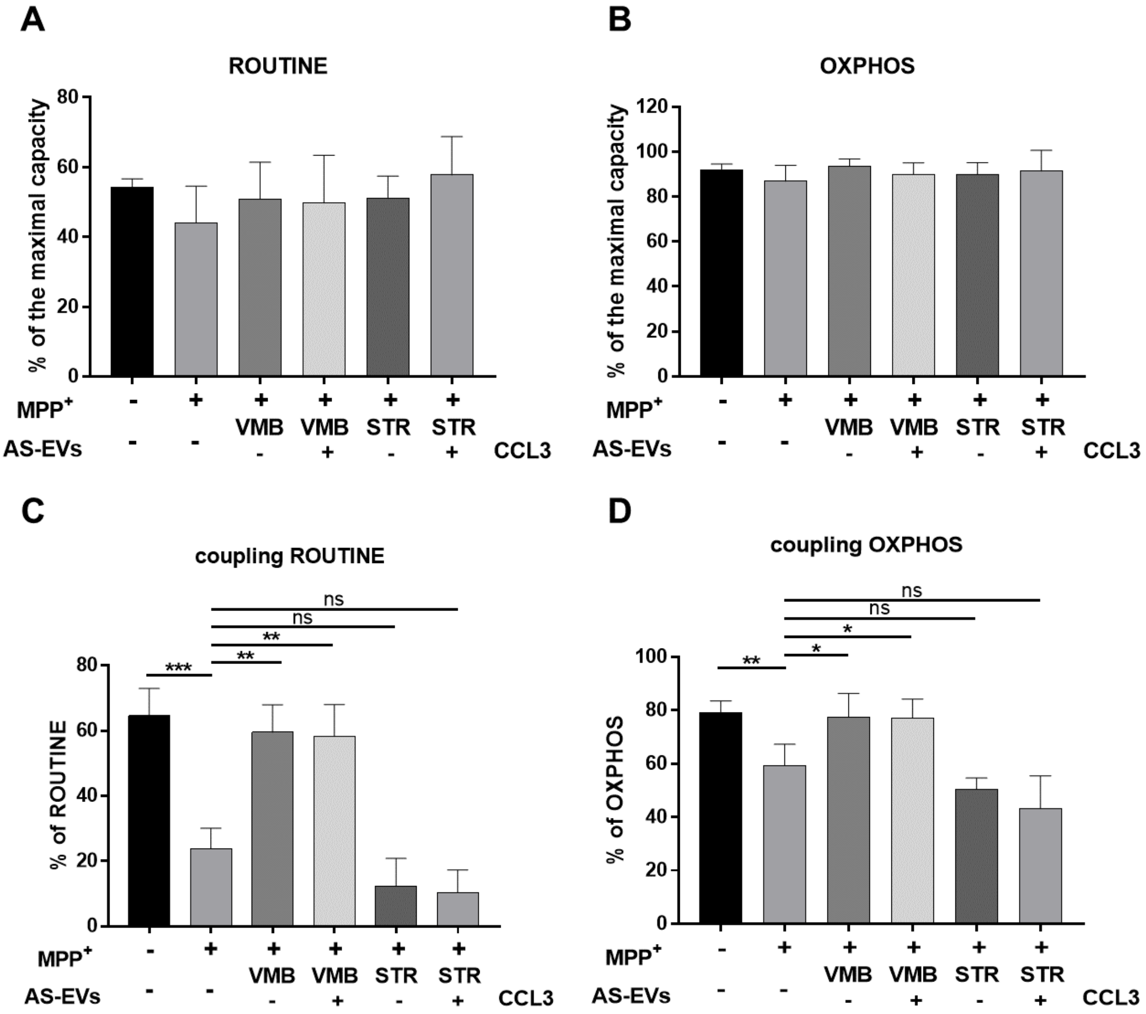


**Figure S5** Analysis of $O_2$ flow coupling efficiency in ROUTINE and OXPHOS upon different conditions. A-B) Analysis of $O_2$ flow expressed as percentage of the maximal ETS capacity calculated in ROUTINE (A) and OXPHOS (B) obtained in intact and permeabilized cells, respectively. As expected, $MPP^+$ does not affect the overall $O_2$ flux in the two states. Additionally, no variation was observed after addition of AS-EVs. C-D) Coupling efficiency of ROUTINE (C) and OXPHOS (D) of SH-SY5Y cells upon different neurotoxin and EVs treatment. According to the ATP production data, only VMB-AS-EVs were able to increase the rate of coupling between the oxidative phosphorylation and ATP production. Data are expressed as percentage of each specific state. Data are expressed as means ± SD. One-way ANOVA with Tukey's multiple comparison. In (C): $**p < 0.01$ ($MPP^+$ vs. $MPP^+$ + VMB-AS-EVs ± CCL3), $***p<0.001$ (CTRL vs. $MPP^+$), ns: not significant. In (d) data were analyzed by t-test $*p < 0.05$ ($MPP^+$ vs. $MPP^+$ + VMB-AS-EVs ± CCL3), $**p < 0.01$ (CTRL vs. $MPP^+$), ns: not significant
